# The CRF-R1 regulation of VTA-GABAergic plasticity is suppressed by CB1 receptor inhibition following chronic exposure to ethanol

**DOI:** 10.1101/272997

**Authors:** Benjamin Harlan, Howard Becker, John Woodward, Arthur Riegel

## Abstract

Dopamine neurons in the ventral tegmental area (VTA) influence learned behaviors and neuropsychiatric diseases including addiction. The stress peptide corticotrophin-releasing factor (CRF) contributes to relapse to drug and alcohol seeking following withdrawal, although the cellular actions are poorly understood. In this study, we show that presynaptic CRF type 1 receptors (CRF-R1) potentiate GABA release onto mouse VTA dopamine neurons via a PKC-Ca^2+^ signaling mechanism. In na1ve animals, activation of CRF-R1 by bath application of CRF or ethanol enhanced GABA_A_ inhibitory postsynaptic currents (IPSCs). Following three days of withdrawal from four weekly cycles of chronic intermittent ethanol (CIE) vapor exposure, spontaneous IPSC frequency was enhanced while CRF- and ethanol-potentiation of IPSCs was intact. However, withdrawal for 3 weeks or more was associated with reduced spontaneous IPSC frequency and diminished CRF and ethanol responses. Long-term withdrawal was also accompanied by decreased sensitivity to the CB1 receptor agonist WIN55212 as well as greatly enhanced sensitivity to the CB1 antagonist AM-251. Inclusion of BAPTA in the internal recording solution restored the responsiveness to CRF or ethanol and reduced the potentiating actions of AM251. Together, these data suggest that GABA_A_ inhibition of VTA dopamine neurons is regulated by presynaptic actions of CRF and endocannabinoids and that long-term withdrawal from CIE treatment enhances endocannabinoid mediated inhibition, thereby suppressing CRF facilitation of GABA release. Such findings have implications for understanding the impact of chronic alcohol on stress-related, dopamine-mediated alcohol seeking behaviors.

## Introduction

Maladaptive changes in the mesocorticolimbic reward system contribute to or exacerbate neuropsychiatric disorders, including depression, anxiety and notably, addiction (for reviews see (Cui *et al*, 2013; Morikawa and Morrisett, 2010)). This system is essential for encoding environmental cues that predict favorable outcomes. Both stress and drugs of abuse (including alcohol) target the reward pathway, that originates from dopamine neurons in the ventral tegmental area (VTA) (Anstrom and Woodward, 2005; Tidey and Miczek, 1996).

The transition from moderate controlled drinking (<2 drinks per day) to heavy drinking (>5 drinks per day) often involves intermittent bouts of binge consumption of ethanol, culminating in dependence (Becker, 2017). Acute administration of ethanol increases the release of mesolimbic dopamine in VTA projection areas (Di Chiara and Imperato, 1988; Yim *et al*, 1998; Zapata and Shippenberg, 2006). Prolonged excessive alcohol consumption is a potent stressor and produces persistent dysregulation of brain reward systems, including the VTA, as well as stress/anti-stress responsivity (Becker, 2017).

Converging evidence shows that the stress hormone corticotropin-releasing factor (CRF) modulates dependence-induced EtOH intake and that CRF signaling itself becomes dysregulated (Pomrenze *et al*, 2017). CRF exerts its effects through two G-protein coupled receptors: Type 1 (CRF-R1) and 2 (CRF-R2) (Bale and Vale, 2004; Burke and Miczek, 2013). Possible sources of CRF include the hypothalamus, BNST and central extended amygdala (CeA) (Rodaros *et al*, 2007; Silberman *et al*, 2013). In the rodent VTA, CRF immunoreactive axon terminals form both asymmetrical (glutamate) and symmetrical (GABAergic) synapses converge onto tyrosine hydroxylase-positive dopamine neurons (Tagliaferro and Morales, 2008). Enhanced CRF-R1-mediated signaling either promotes, or is at least required, in the VTA for excessive ethanol consumption in rodents (Hwa *et al*, 2013; Sparta *et al*, 2013). Intra-VTA infusions of a CRF-R1 antagonist decreased intermittent ethanol intake in stressed and non-stressed mice, but interestingly not in mice given continuous access to ethanol (Hwa *et al*, 2013). Although the VTA CRF-R1 action coincides temporally with repeated withdrawal, CRF-R2 likely also plays a role (Rinker *et al*, 2017). The CRF-R1 circuit between CeA and VTA may collectively promote binge alcohol intake during dependence (Rinker *et al*, 2017). Systemic administration of CRF-R1 antagonists that reduce operant self-administration in rats also reduce the withdrawal-induced enhancement of GABA release when infused locally into the CeA (Roberto *et al*, 2010a; 2010b). This GABA reduction in the CeA has been associated with negative reinforcement mechanisms (Roberto *et al*, 2010b). In the VTA, activation of GABA_A_ receptors are known to profoundly limit dopamine cell bursting (Overton and Clark, 1997) and conversely, suppression of GABA_A_ transmission switches dopaminergic neurons into a burst firing pattern (Paladini and Tepper, 1999; Paladini *et al*, 1999; Tepper *et al*, 1995). However, the regulation of GABA release in the VTA by CRF-R1 before or during alcohol dependence is, by comparison to the CeA, less well understood.

Understanding the interaction between CRF-R1 and EtOH on GABA VTA terminals is also interesting considering the link between ethanol and the endocannabinoid (CB1 receptor) regulation of VTA GABA release. Acute administration of ethanol (Di Chiara and Imperato, 1988; Yim *et al*, 1998) or a CB1 agonist (Cheer *et al*, 2004) enhances the activity of VTA dopamine neurons and dopamine levels in the nucleus accumbens (Gessa *et al*, 1998; Tanda *et al*, 1997). Conversely, CB1 receptor antagonists decrease both the ethanol related voluntary intake in rodents (Arnone *et al*, 1997; Colombo *et al*, 1998; Freedland *et al*, 2001) and the increases in accumbal dopamine (Cheer *et al*, 2007). Such ethanol actions are absent in CB1 knockout mice (Hungund *et al*, 2003). Because the acute withdrawal from ethanol robustly elevates endocannabinoid production (Basavarajappa *et al*, 2003), the role of CRF-R1 on GABA transmission (and by association VTA output) may be altered during alcohol dependence.

Here we explore the possible mechanistic link between CRF-R1 and GABA and determine if ethanol dependence alters this relationship through an action of endocannabinoids. First, we examined the acute actions of CRF on GABA release in the VTA using whole-cell patch-clamp recordings in brain tissue of na1ve mice. Second, we explored this interaction in mice exposed to repeated cycles of chronic intermittent ethanol (CIE) vapor exposure. The CIE protocol is associated with recurring surges of CRF and is an established rodent model of alcohol dependence (Becker and Lopez, 2004). We identify a presynaptic CRF-R1 mediated mechanism involving PKC and intracellular calcium stores that facilitates GABA release onto VTA DA neurons. We show that in the weeks following CIE exposure, CRF’s action is suppressed by a persistent functional enhancement of endocannabinoid mediated CB1 receptor inhibition at VTA GABA synapses. Together, these data demonstrate an enduring adaptation that is distinct from the classic somatic withdrawal symptoms that typically recede in 48 hours (Diana *et al*, 1996). This persistent adaptation in CRF-mediated control of VTA-GABA synapses may play a key role in alcohol addiction.

## Materials and Methods

### Slices and Solutions

All experiments were conducted according to the National Institutes of Health Guidelines for the Care and Use of Laboratory Animals. Brain slices were prepared as described previously (Williams et al., 1984). Briefly, horizontal slices (220 µm) of the ventral mid-brain were prepared from adult (P60-P160) male C57BL-6J mice. Slices were cut using a Vibratome (Leica) in ice-cold ACSF containing 126 mM NaCl, 2.5 mM KCl, 1.2 mM MgCl_2_, 2.4 mM CaCl_2_, 1.4 mM NaH_2_PO_4_, 25 mM NaHCO_3_, 11 mM D-glucose and 0.4 mM ascorbate, and MK-801 (0.01 mM). Slices were stored in oxygenated (95% O_2_%-5% CO_2_) ACSF containing MK-801 (0.01 mM) for 30 min at 33C before recording. Electrophysiological recordings were performed at 33^°^C. The perfusion rate of oxygenated ACSF (95% O_2_%-5% CO_2_) was 2 ml/min. Neurons were visualized with an Olympus BX51WI (Olympus America) microscope.

### Whole-Cell Recordings

Recordings were performed dopamine neurons in brain slices from adult (P60-P160) male C57BL-6J mice similar to that described previously (Williams *et al*, 2014). Briefly, whole-cell recordings were made using Axopatch 700B amplifier (Axon Instruments). Recordings were collected using AxoGraph X (AxoGraph Scientific, Sydney, Australia), filtered at 1-2 kHz and digitized at 2-5 kHz. Neurons were voltage clamped at −60 mV using 1.5-2.0 MO pipettes. Pipette internal solution contained 57 KCl, 57 K-methylsulphate, 20 mM NaCl, 1.5mM MgCl_2_, 5 mM HEPES, 2 mM ATP, 0.3 mM GTP, and 10 phosphocreatine (pH 7.3; 265-270 mOsm). Unless noted, the internal solution also contained 0.1 mM EGTA. Series resistance (3-10 MO) was compensated at 80%. All recordings were performed in dopamine neurons identified by their large cell bodies (20 µm), the characteristic pacemaker-like firing (1-5 Hz) observed in the cell-attached mode, and the presence of a large (> 300 pA) hyperpolarization-induced */*_h_ current. These cells also display a hyperpolarizing mGluR-mediated current carried by the sK channel that is observed only in dopamine neurons in midbrain slices (Marino *et al*, 2001). Cells were visualized using a 60x water-immersion objective on an upright microscope (Olympus America). To evoke inhibitory postsynaptic currents (IPSCs), a bipolar stimulating electrode was placed such that the cell of interest resided between the two ends of the stimulating electrode. A stimulation intensity was selected to evoke an amplitude of 30% of maximal IPSC amplitude. A paired pulse protocol (0.2 ms duration, 50 ms interval, 0.06 Hz) was used to evoke IPSCs mediated by the activation of GABA_A_ receptors. The GABA_A_ IPSCs were isolated pharmacologically with NBQX (3 µM), SCH-23390 (1 µM), CGP-55845 (10 µM), and sulpiride (200 nM). Spontaneous miniature IPSCs (mIPSCs) were recorded in the presence of tetrodotoxin (TTX; 200 nM). The sIPSCs and mIPSC amplitudes and inter-event interval times were measured with AxographX using a sliding algorithm template. For blockade of CRF receptors, CRF-R1/R2 antagonists were incubated for 10-15 min prior to application of CRF.

### Drugs

Drugs were applied by extracellular perfusion. Corticotropin Releasing Factor (CRF) and the urocortin drugs (UCN, UCN-2, and UCN-3) were obtained from American Peptide (Sunnyvale, CA). CGP-55845, chelerythrine, PdBU, sulpiride, K41498, and CP154156 were obtained from Tocris Bioscience (Minneapolis, MN). Astressin 2B, H-89, TTX, CPA, picrotoxin, SCH-23390, WIN55212, NPY, NBQX, U69596, and AM251 were obtained from Ascent Scientific (now Abcam; Cambridge, MA). An ethanol concentration (50 mM; Sigma Aldrich) was selected for its relevance to previous work (Roberto and Varodayan, 2017).

### Ethanol Treatment

All experiments using ethanol- and air-treated mice were performed in age-matched animals. Treated animals were subjected to a chronic intermittent ethanol (CIE) exposure model as described previously (Becker and Lopez, 2004; Lopez and Becker, 2005; Dhaher et al., 2008; Griffin et al., 2009a; 2009b; Lopez et al., 2012). Briefly, one group of mice (ethanol [CIE] group) received pyrazole (1 mmol/kg) plus a loading dose of ethanol (3 g/kg) immediately prior to entering the CIE chamber each day during four exposures (16 h/d, 4 days on, and 72 hrs off) of ethanol exposure in custom inhalation chambers. Ethanol vapor concentrations were monitored daily to ensure that the inhalation conditions produced stable blood ethanol levels (BEC) between 175 and 225 mg/dl (Griffin *et al*, 2009). BEC was assessed once each week in sentinel mice by sampling blood from the retro-orbital sinus immediately upon removal from the chamber. Air-exposed mice (control group) were given a dose of pyrazole before entering the cambers and were exposed to control (air) inhalation conditions. After four weeks of treatment, all mice were removed from the vapor chambers and returned to their home cage. The repeated exposure to ethanol results in somatic withdrawal symptoms that typically recede in ∼48 h (Diana *et al*, 1996). To differentiate between withdrawal and other potentially more enduring mechanisms, we operationally defined the period following cessation of CIE exposure as experimenter-induced withdrawal. Mice were killed 3 (short-term withdrawal) or 25-45 days (long-term withdrawal) following the final ethanol or air exposure for tissue collection.

### Statistical Analysis

Data were collected and later analyzed offline on a Macintosh-mini computer (Apple, Sunnyvale, CA) using AxoGraph X (AxoGraph X 1.6.4) and Chart 5.2.2 (AD Instruments). Data are presented throughout as the mean ± SEM. Statistical analyses were performed using Prism (GraphPad Software, La Jolla, CA). Paired and unpaired *t*-tests were used where necessary for statistical comparisons. Differences were considered significant if *P*<0.05. One-way ANOVAs or two-way ANOVAs followed by Dunnett’s *post hoc* test were used to determine the effects of multiple treatments.

## Results

### CRF potentiates GABA_A_ receptor mediated IPSCs via presynaptic CRF-R1

We recorded stimulus-evoked GABA_A_ mediated inhibitory post-synaptic currents (eIPSCs) from VTA dopamine neurons in acute brain slices from C57BL-6J mice. The concentration dependence of two non-selective agonists (CRF and UCN) and one CRF-R2 agonist (UCN-3) were evaluated after 10-minute bath application, over a range of 10 - 300 nM (Figure 1a). Both CRF and UCN potentiated the amplitude of eIPSCs with near maximal effects at 200 nM (Figure 1a; CRF: t_(12)_ = 20.19, p < 0.0001 relative to baseline; UCN: t_(10)_ = 26.01, p < 0.0001 relative to baseline). UCN-3 had no measurable effect on eIPSCs (Figure 1a, 200 nM UCN-3: t_(8)_ = 1.148, p = 0.1884 relative to baseline). The EC_50_ for CRF and UCN was 34 nM, 41 nM, respectively (Figure 1a). At 300 nM, CRF also caused a persistent inward current (8.1 ± 3.1 pA), consistent with previous reports (Beckstead *et al*, 2009a). At the end of the experiments, the GABA_A_ receptor antagonist picrotoxin (picro, 100 µM) was applied to confirm the specificity of GABA_A_ receptor mediated currents (Figure 1a).

**Figure 1.**
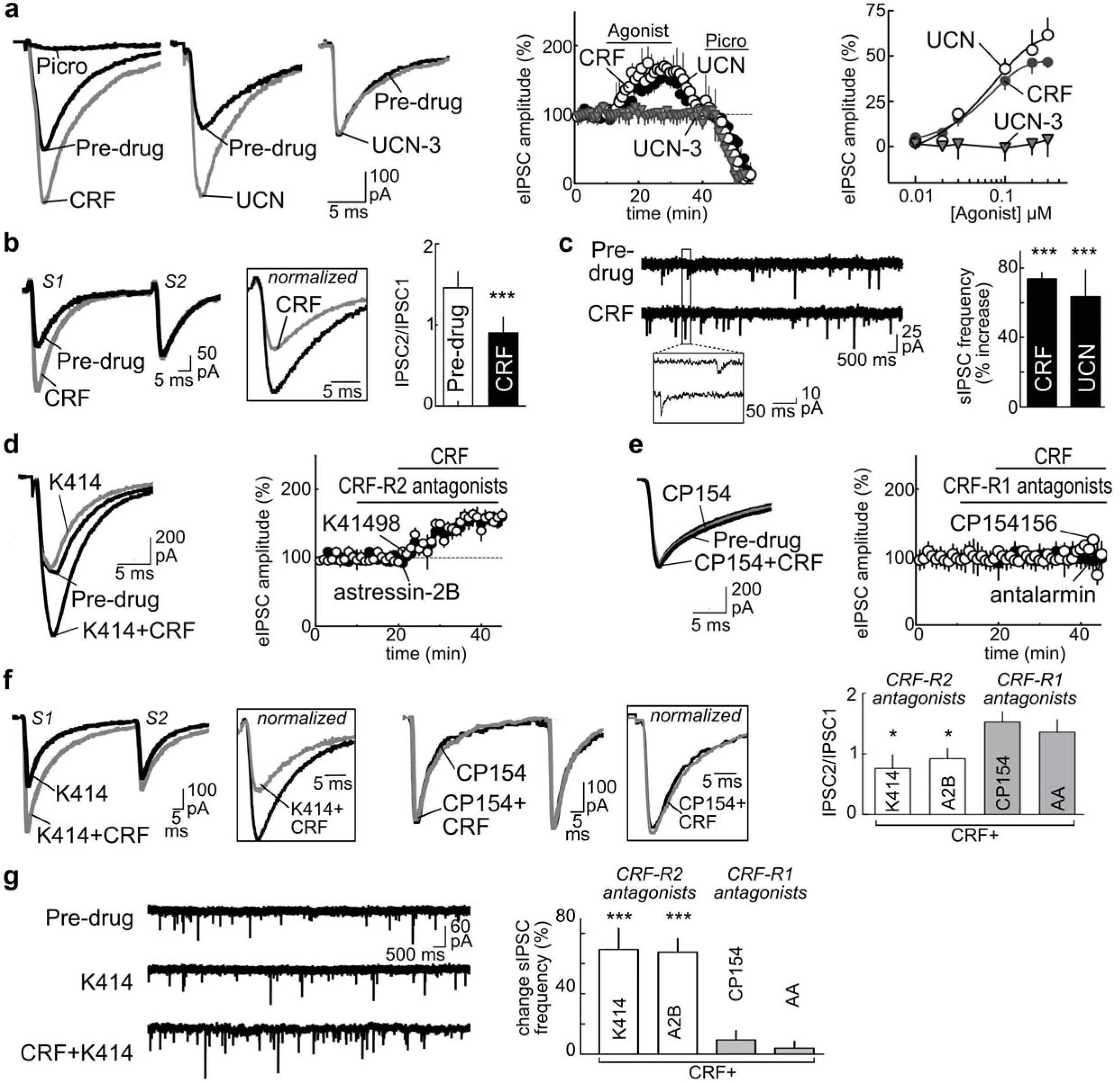
Presynaptic CRF-1 receptors potentiate GABA_A_ IPSCs in VTA DA neurons. (**a**) Traces of stimulus-evoked, picrotoxin-sensitive IPSCs showing potentiation by CRF (200 nM) and urocortin (UCN; 200 nM), but not the CRF-R2 agonist UCN-3 (300 nM). For the timeline, the amplitude of the evoked current was normalized for each cell using the mean amplitude recorded during the first 5 min and plotted as function of time (% of baseline, mean ± SEM). Although CRF (200 nM) and UCN (200 nM) potentiated IPSCs, UCN-3 did not. The smooth curves in the agonist concentration-response plot are the best fit to the data by the logistic equation. (**b**) CRF (300 nM) increased the amplitude of the first eIPSC (S1), with little measurable change in the second eIPSC (S2). *Inset* shows the first eIPSCs (S1) normalized to the second eIPSC (S2). CRF significantly decreased the paired pulse ratio (PPR; IPSC2/IPSC1, ^***^ p<0.001. (**c**) CRF (300 nM) enhances the frequency of spontaneous IPSCs (sIPSCs). *Inset* shows magnification of sIPSCs. Both UCN and CRF increased the frequency of sIPCS relative to their baseline (^***^ p<0.001). (**d, e**) The CRF (300 nM) potentiation is blocked by co-application of (e) CRF-R1 antagonists (CP154156 (CP154) or antalarmin (AA), but not (d) CRF-R2 antagonists (K41498 (K414) or astressin-2B (A2B) (^*^ p < 0.05). (**f**) The CRF (300 nM) shift in the PPR is reduced by CRF-R1 blockers (CP154 or AA), but not CRF-R2 blockers (K414 or A2B). One-way ANOVA, ^*^ p < 0.05 vs CRF alone (Dunnett’s post test). (**g**) The CRF (200 nM) enhancement of sIPSC frequency is prevented by co-application of CRF-R1 blockers (CP154 or AA), but not CRF-R2 blockers (K414 or A2B). One-way ANOVA, ^***^ p < 0.001 vs CRF alone (Dunnett’s post test). (a-g) Number of cells/mice per group: (a) CRF 13/7; UCN 11/6, UCN-3 9/4 for both timelines and concentration-response curves; (b) 7/4; (c) 7/5, (d and e) K414 9/5, A2B 9/5, CP154 8/4, AA 7/3; (f) 7/3-4 per group; (g) CRF+K414 7/4, and the following groups contained 8-9/4-5: UCN+A2B, CRF+A2B, CRF+CP154, and CRF+AA.

To assess whether the CRF acted at a pre- or post-synaptic site, we recorded paired pulse ratios (PPRs, IPSC2/IPSC1, Figure 1b). PPRs typically correlate inversely with release probability (Debanne *et al*, 1996). CRF (300 nM) increased the amplitude of the first eIPSC (Figure 1b, S1, paired t-test t_(6)_ = 9.625, p < 0.0001), with little measurable change in the second eIPSC (Figure 1b, S2, paired t-test t_(6)_ = 0.7661, p = 0.2663). This resulted in a significant decrease in the PPR in the presence of CRF (Figure 1b *inset*, paired t-test, t_(6)_ = 8.2601, p = 0.0002), thus suggesting a presynaptic locus for CRF’s actions on GABA_A_ eIPSCs.

While measures of PPR are commonly used to assess changes in presynaptic release, other modifications of short-term plasticity can alter the PPR independent of changes in release probability (Trussell *et al*, 1993). To confirm a presynaptic locus of CRF action, we measured the frequency and amplitude of spontaneous and miniature IPSCs (sIPSCs and mIPSCs, respectively). CRF or UCN increased the sIPSC frequency in all cells tested (Figure 1c; CRF: t_(7)_ = 4.567, p = 0.0003; UCN: t_(7)_ = 4.324, p = 0.0003). Similar increases were observed with CRF on mIPSC recordings (medium containing 200 nM TTX (t_(7)_ = 5.885, p = 0.0023). CRF did not significantly change the amplitude of sIPSCs or mIPSCs (sIPSCs: 5.56% ± 4.6%, t_(14)_ = 0.03597, p = 0.4860; mIPSCs: 2.94% ± 11.13%, t_(7)_ = 0.5172, p = 0.3105). Together, the results from the PPR studies and sIPSC/mIPSC recordings are consistent with a presynaptic action of CRF on GABA synapses on to VTA DA neurons.

### CRF potentiation of GABAA receptor mediated IPSCs requires CRF-R1

To assess whether the CRF facilitation of GABA_A_ currents was mediated by CRF-R1 or -R2, CRF was applied in the presence of blockers for CRF-R1 (300 nM CP154156, CP154; 200 nM antalarmin, AA) or CRF-R2 (300 nM K41498, K414; 200 nM astressin-2B, A2B). Slices were incubated with antagonists 10 min prior to and during CRF challenges. The resulting changes in evoked currents, the PPR, and the frequency of sIPSCs was examined.

Neither of the CRF-R2 blockers, K414 or A2B, prevented the CRF-potentiation of eIPSCs (Figure 1d; CRF+K414: t_(6)_ = 3.564, p = 0.0121; CRF+A2B: t_(7)_ = 4.234, p = 0.0223). However, co-application of the CRF-R1 antagonists CP154 or AA completely blocked the CRF increase of eIPSCs (Figure 1e; CRF+CP154 t_(5)_ = 0.03185, p = 0.5241; CRF+AA: t_(7)_ = 0.02354, p = 0.7123). To verify that these effects were mediated by presynaptic CRF-R1, we evaluated the PPR (Figure 1f) and sIPSC frequency (Figure 1g) in the presence of selective CRF receptor antagonists. A one-way between subject ANOVA evaluating the effects of the four antagonists on the CRF mediated changes in the PPR (Figure 1f) or in the frequency of sIPSC (Figure 1g) indicated significant differences at the p<0.05 level (PPR *F*_(4,22)_= 3.884, p = 0.0156; frequency *F*_(4,64)_ *=* 13.16, p < 0.0001). Separate post hoc tests (multiple comparisons Dunnett’s) revealed that the CRF changes in the PPR (Figure 1f) were absent when blocking CRF-R1 with CP154 (p = 0.9997) or AA (p = 0.9745), but were observed during co-application of the CRF-R2 blockers K414 (p = 0.0137) or A2B (p = 0.0413). Post hoc tests revealed that the CRF changes in frequency (Figure 1g) followed the same pattern: CRF changes were absent when blocking CRF-R1 with CP154 (p = 0.9831) or AA (p = 0.7991), but observed during co-application of the CRF-R2 blockers K414 (p < 0.0001) or A2B (p < 0.001). This indicates that the CRF related changes in GABA release are mediated by presynaptic CRF-R1 receptors.

The possibility that endogenous release of CRF tonically activates CRF receptors was also tested using selective antagonists. Application of either of the CRF-R1 or CRF-R2 antagonists alone, without CRF, produced no measurable change in eIPSCs (one way ANOVA, *F*_(4,41)_= 0.3584, p = 0.8367), nor in the frequency of sIPSCs (one way ANOVA F_(4, 30)_ = 0.0194, p = 0.9618), suggesting the absence of a measurable tone on CRF receptors in slices prepared from na1ve mice. Together, the results from the PPR studies and sIPSC/mIPSC recordings are consistent with a presynaptic action of CRF via CRF-R1 on GABA synapses on to VTA DA neurons.

### CRF-R1 acts through Protein Kinase C and calcium induced calcium release

CRF receptors couple to the Gs alpha subunit to activate adenylyl cyclase, as well as the Gq alpha subunit to activate PLC-PKC (Arzt and Holsboer, 2006). To better understand the intracellular pathway regulating the CRF potentiation of GABA onto DA cells, we evaluated eIPSCs during co-manipulation of CRF-R1 and the PKC enzyme.

As previously shown, CRF increased the amplitude of eIPSCs (Figure 2a; t_(9)_ = 4.6327, p = 0.0143) and these responses were further potentiated by phorbol-ester-dybutyrate (PdBU), an activator of PKC (Figure 2a; t_(7)_ = 4.1032, p = 0.0178). However, if PdBU (1 µM) was applied first, application of CRF (300 nM) produced no additional increase in eIPSC amplitude (Figures 2b). The lack of a CRF action in the presence of PdBU was not due to a ceiling effect of maximal chloride conductance, because at the end of the experiment a more intense synaptic stimulation (∼35%) further increased the inward current (158.4% ± 51.6% relative to baseline, t_(2)_ = 16.99, p = 0.0017). The PdBU-potentiation of eIPSC amplitude was not reduced by the CRF-R1 antagonist K41498 (PdBU vs PdBu+K414: 0.56% ± 11.7%, t_(5)_ = 0.3577, p < 0.3676), suggesting an activation of PKC activity downstream of CRF-R1.

**Figure 2.**
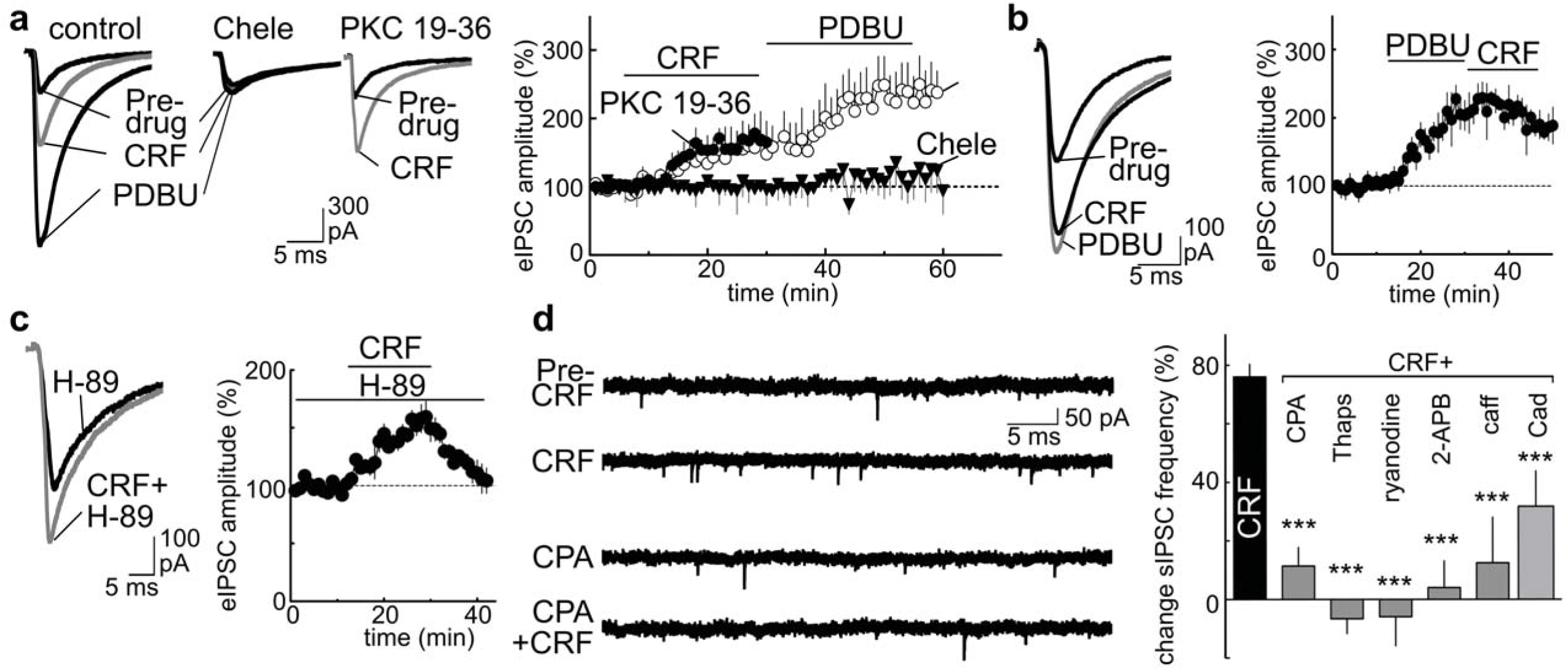
The CRF potentiation of GABA_A_ receptor IPSCs requires PKC and intracellular calcium stores. (**a**) The CRF (200 nM) potentiation of eIPSCs is further increased by the PKC activator Phorbol 12,13-dibutyrate (PdBU; 1 µM). Bath incubation of slices with the PKC inhibitor chelerythrine (Chele, 1 µM), blocked the actions of CRF (200 nM) and PDBU (1 µM), whereas inclusion of the cell impermeable PKC inhibitor PKC19-36 (300 nM) in the internal recording solution did not. The timeline shows the eIPSC amplitude as a percentage of pre-drug baseline ± SEM, normalized for each cell using the mean amplitude recorded during the first 5 min of pre-drug application and plotted as a function of time. (**b**) Application of PdBU (1 µM) prior to CRF (200 nM) reduces the CRF potentiation of eIPSCs. (**c**) Incubation of brain slices with the PKA inhibitor (H-89: 10 µM; incubation >30 min) did not prevent the CRF-potentiation of eIPSCs. (**d**) Summary bars showing pharmacological manipulation of calcium stores reduces the actions of CRF. Drugs include antagonists/inhibitors of: Ca^2+^-ATPases [cyclopiazonic acid (CPA 10 µM), thapsigargin (Thaps, 5 µM)], ryanodine receptors (ryanodine, 50 µM), IP3 receptors (2-aminoethoxydiphenyl-borate (2-APB, 10 µM)); voltage-dependent calcium channels (cadmium, Cad 300 µM). Also tested is caffeine (caff, 10 mM), an agonist of ryanodine receptors used to deplete intracellular calcium stores. CRF (300 nM) data in bar graph is duplicate from Figure 1c, shown again for comparison. One-way ANOVA, ^***^ p < 0.001 vs CRF alone (Dunnett’s post test). (a-d) Number of cells/mice per group: (a) control 10/5, Chele 15/7, PKC 19-36 8/4; (b) 6/3; (c) 10/5; (d) CRF 15/7, CRF+CPA 8/4, CRF+Thaps 5/3, CRF+ryanodine 6/4, CRF + 2-APB 5/3, CRF+caff, 6/3, Cad 8/4.

To confirm whether PKC is involved in this response, we performed similar experiments in the presence of various PKC inhibitors. Treatment of slices with chelerythrine (chele: 1 µM, >30 min), which inhibits the catalytic subunit of PKC (Herbert *et al*, 1990; Ko *et al*, 1990), blocked the effects of CRF or PdBU on eIPSCs (Figures 2a; CRF+chele t_(7)_ = 0.03016, p = 0.5781). Since chelerythrine is membrane permeable (Herbert *et al*, 1990) and will block both presynaptic and postsynaptic PKC enzymes, additional experiments were performed using an internal solution containing the non-membrane permeable pseudo-substrate blocker of PKC, PKC 19-36 (5 µM internal solution; infusion >20 min) (Yasunari *et al*, 1996). In cells loaded with PKC 19-36, CRF still potentiated evoked IPSCs, consistent with a presynaptic locus of CRF action (Figures 2a; t_(5)_ = 4.2340, p = 0.0117). In contrast to the effects observed with PKC inhibitors, CRF-potentiation of IPSCs was unaffected by the PKA inhibitor H89 (10 µM; >30 min incubation; present during recording) (Figure 2c; t_(4)_ = 4.0124, p = 0.0223). This CRF-R1/PKC-dependent action in mice differs from the CRF-R2/PKA-dependent mechanism we previously observed in rats (Williams *et al*, 2014). Thus, CRF and PdBU enhance GABA_A_ IPSCs in mice via a presynaptic PKC-dependent mechanism.

Neuropeptides can regulate transmitter release by a kinase-dependent mobilization of calcium release from intracellular stores (Arzt and Holsboer, 2006). Accordingly, we observed that depletion of stores by incubating slices (> 15 min) with cyclopiazonic acid (CPA; 10 µM) reduced the CRF facilitation of mIPSCs frequency (29.2% ± 2.84 % decrease relative to pre-CPA; t_(9)_ = 10.89, p < 0.0001), but not the amplitude (mIPSC amplitudes 9.26% ± 13.12% decrease relative to baseline, t_(9)_ = 0.5246, p = 0.3063). CPA and TTX treatments also decreased the frequency of recordable mIPSCs. However, we confirmed the results by measuring sIPSCs, which have a higher basal frequency than mIPSCs. Slices were treated with CPA, thapsigargin (5 µM), ryanodine, caffeine or 2-APB (Figure 2d)-drugs that either block IP_3_/ryanodine receptors or deplete calcium stores (Maruyama *et al*, 1997; Morikawa *et al*, 2000; Verkhratsky, 2005). Because the frequency of sIPSCs varied considerably between slices/cells even under control conditions without antagonist (range 4-8Hz, mean 4.39 ± 0.43Hz), we determined the CRF-related increase in the frequency within cells before CRF (in the presence of the antagonist) and during CRF. A one-way between subject ANOVA comparing CRF responses in the presence of the antagonists indicated significant differences (Figures 2d; *F*_(6,45)_16.21, p <0.0001). Post hoc tests (multiple comparisons Dunnett’s) indicated the change was significant for all the compounds (Figures 2d; p <0.0001 all drugs, except cadmium; cadmium, p <0.006). The observation that these different treatments all reduced the CRF action on sIPSCs (Figure 2d), indicates a need for functional calcium stores. It was notable that treatment with the non-selective calcium channel blocker cadmium reduced, but did not block, the CRF action on sIPSCs (Figure 2d). Thus, although CRF facilitation does not require extracellular calcium, calcium entering through voltage gated calcium channels likely potentiates the calcium release from intracellular stores (calcium-induced calcium-release)(Verkhratsky, 2005).

### Reduced CRF and EtOH effects on GABA_A_ currents after long-term withdrawal from chronic intermittent ethanol exposure (CIE)

Similar to other brain regions, ethanol can enhance GABA release onto VTA dopamine cells (Theile *et al*, 2008; Zhu and Lovinger, 2006). Using VTA slices from na1ve mice, we measured the frequency of GABA_A_ IPSCs during ethanol application (50 mM; 10 min) and confirmed that the potentiation also occurs under our recording conditions (Figure 3a; t_(5)_ = 5.733, p = 0.0023). The ethanol action may require CRF-R1 and the release of calcium from intracellular stores, as observed in other brain regions (Roberto and Varodayan, 2017), so we wondered if this also occurs in the VTA. To evaluate this, we measured the ethanol enhancement of sIPSCs frequency in slices from na1ve mice during bath application of: 1) ethanol, 2) ethanol plus the CRF-R1 antagonist CP154, or 3) ethanol plus CRF. A one-way ANOVA comparing responses under these conditions indicated no significant differences (Figures 3a; *F*_(2,32)_*=* 0.1467, p = 0.8644). The absence of an additive effect of CRF on ethanol or any reduction in the ethanol action by the CRF-R1 antagonist suggests that the “stress-like” actions of ethanol may reflect intracellular alterations somewhere downstream of the CRF-R1 receptor.

**Figure 3.**
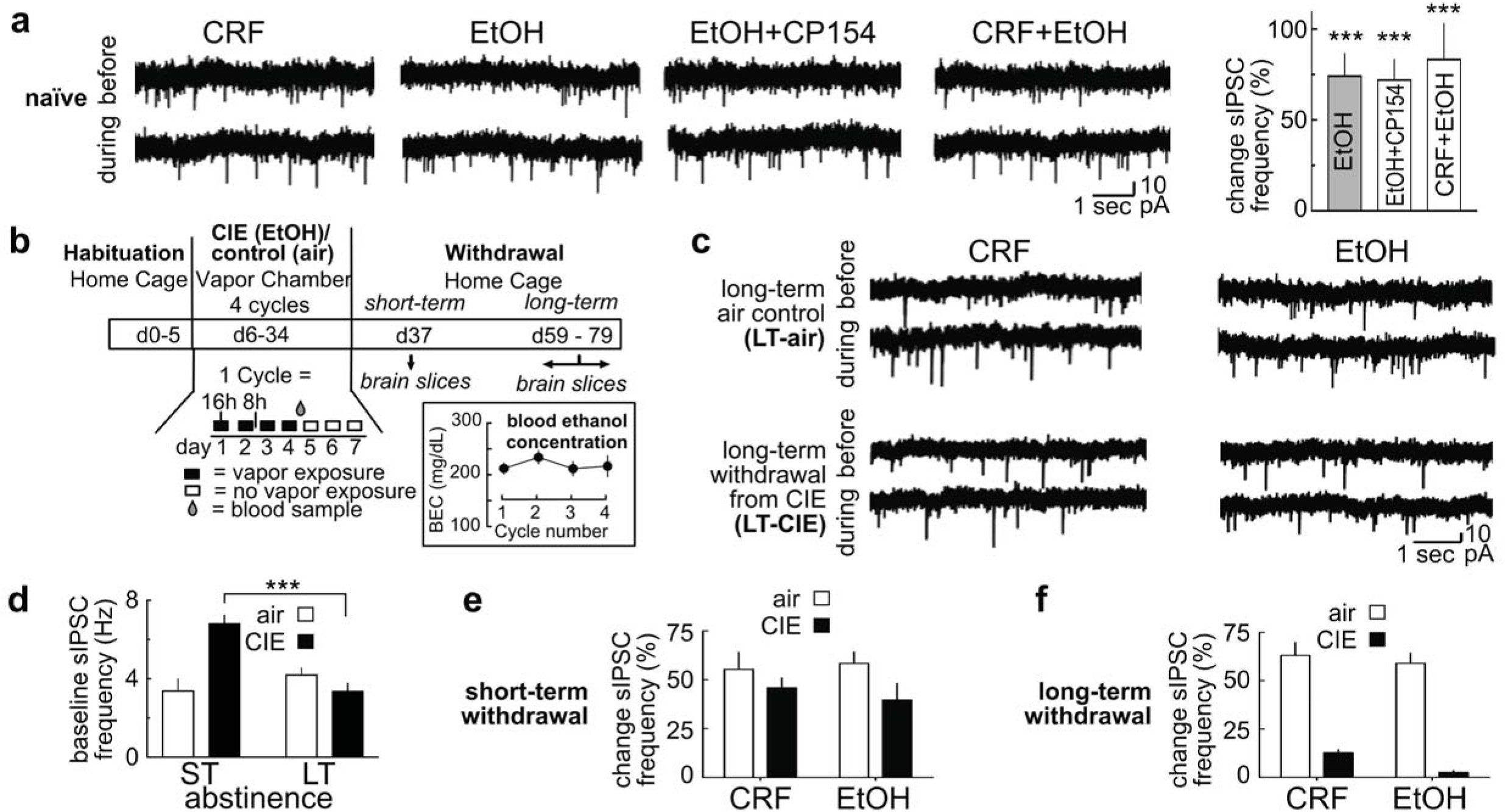
Reduction in responses to CRF or ethanol following long-term withdrawal from chronic intermittent ethanol (CIE) exposure. (**a**) In neurons from na1ve mice, bath application of CRF (300 nM) or ethanol (EtOH; 50 mM) increases the sIPSC frequency relative to their baseline to a similar extent. Ethanol responses are not significantly altered by co-application of the CRF-R1 antagonist CP154156 (CP154; 300 nM; incubation>12 min) or CRF (CRF + EtOH). The CRF traces are shown for comparison. See Figure 1c for average CRF data. ^***^ p<0.001 compared to baseline. (**b**) Schematic showing experimental timeline for chronic intermittent ethanol (CIE) exposure. Habituation (5 days) was followed by vapor chamber exposure to ethanol (or air for controls) over 4 cycles (16 hours per day; 4 days per week). Blood ethanol concentrations (BEC) were measured after each cycle and averaged ∼210 mg/dl-a concentration sufficient to produce tolerance and dependence (see Methods). After the final air/CIE treatment, mice were returned to the home cage. Brain slices were later prepared to compare physiological responses after a period of short-term withdrawal (ST; 3d after treatment) or long-term withdrawal (LT; pooled responses from 20d, 30d and 45d after treatment and reported as a group). (**c**) Sample sIPSC traces from LT-air (top) and LT-CIE (bottom) cells before (baseline) and during bath application of CRF (300 nM) or ethanol (EtOH; 50 mM). (**d**) Summary graph showing differences in baseline frequency of sIPSC in ST and LT withdrawal groups without application of CRF or ethanol. One-way ANOVA, ^***^ p < 0.001 ST-CIE vs LT-CIE (Dunnett’s post test). (**e**) During ST withdrawal, CRF (200 nM) and ethanol (EtOH; 50 mM) enhance sIPSC frequency similar to control levels. (**f**) During LT-withdrawal, the CRF (200 nM) and EtOH (50 mM) action is reduced in CIE group relative to controls. Two-way ANOVA, p < 0.0001 for effect of treatment. (a-f): Number of cells/mice per group: (a) EtOH 6/4, CP154 + EtOH 6/4, CRF+EtOH 7/5; (d) ST-air 5/4; ST-CIE 6/5, LT-air 5/4, LT-CIE 11/6; (e) ST-air-CRF 18/9; ST-CIE-CRF 13/6, ST-air-EtOH 10/5, ST-CIE-EtOH 17/8. (f) LT-air-CRF 18/9; LT-CIE-CRF 11/6, LT-air-EtOH 10/7, LT-CIE-EtOH 9/6.

Chronic intermittent ethanol (CIE) exposure produces dependence and robustly activates CRF stress systems (Becker and Lopez, 2004; Griffin *et al*, 2009; Lopez and Becker, 2005). So, we predicted that CIE treatment (Figure 3b) would also alter the GABA_A_ responses to CRF and ethanol. To test this, we prepared VTA slices from mice to evaluate air or CIE treatments under two conditions: short-term (ST) withdrawal of 72 hrs or a long-term (LT) withdrawal of 20-45 days. We recorded basal frequency of sIPSCs (in Hz) in the ST and LT groups (Figure 3c and d) and found a significant interaction between treatment and time (two-way ANOVA, F_(1, 21)_ = 4.457, p = 0.0469; post hoc Dunnett’s multiple comparison test: ST-vs LT-CIE p = 0.0007, ST-vs LT-air p = 0.7219). This suggests that the CIE induced change in basal release of GABA is temporary: During the ST withdrawal periods, the frequency of sIPSCs increase, whereas during LT withdrawal periods, the spontaneous release of GABA decreases toward control levels.

It was not clear whether the decrease in sIPSCs represented an active change or simply recovery from the acute effects of CIE. On this basis, we examined the actions of CRF and ethanol on sIPSCs starting with the ST withdrawal group. A two-way ANOVA revealed no significant differences in response to CRF or ethanol for sIPSCs (Figure 3e: treatment (*F*_(1,20)_ = 3.435, p = 0.0786), drug (*F*_(1,20)_ = 0.0016, p = 0.9682); interaction of treatment x drug (*F*_(1,20)_ = 1.14, p = 0.2984)), nor for eIPSCs (treatment (*F*_(1,20)_ = 0.3054, p = 0.5866), drug (*F*_(1,20)_ = 0.1761, p = 0.6792); interaction of treatment x drug (*F*_(1,20)_ = 0.0026, p = 0.9625)). This suggests that during ST withdrawal, the mechanisms that facilitate GABA release remains functional. However, a similar comparison after LT withdrawal did reveal differences in the ability of CRF and ethanol to alter the frequency of sIPSCs (Figure 3c and f). For sIPSCs, a two-way ANOVA showed a significant effect of treatment (*F*_(1,17)_ = 142.2, p < 0.0001), but not drug (*F*_(1,17)_ = 0.8148, p = 0.3793) or interaction (*F*_(1,17)_ = 1.15, p = 0.2986). As a point of control, additional experiments were performed measuring evoked IPSCs and a two-way ANOVA of evoked IPSCs also revealed an effect of treatment (*F*_(1,17)_ = 10.64, p < 0.0046), but not drug (*F*_(1,17)_ = 0.03609, p = 0.8516 or interaction (*F*_(1,17)_ = 0.1804, p = 0.6764). Thus, the actions of CRF and alcohol on the release of GABA appear similar, but measurably change following LT withdrawal from CIE treatments.

### CB1 mediated inhibition opposes the actions of CRF

To determine the signaling mechanism underlying the blunted CRF and ethanol modulation of GABA_A_ currents after 20-45 days of withdrawal from CIE treatment, we retested the response to CRF and the PKC activator PdBU using eIPSCs, as they are easily discernable due to their large size. While CRF and PdBU responses in LT-air mice were robust (Figure 4a; CRF: t_(6)_ = 3.3810, p = 0.0221, PdBU: t_(4)_= 03.6542, p = 0.0323) and resembled those in na1ve tissue as noted previously, no increase was observed in response to PdBU in slices from LT-CIE mice (Figure 4a; CRF: t_(6)_ = 0.3415, p = 0.3524, PdBU: t_(5)_= 0.2153, p = 0.2972). Although acute somatic withdrawal from alcohol is known to increase the synthesis of CRF (Merlo-Pich *et al*, 1995), no evidence of a ceiling effect, nor a CRF tone, was observed, as application of the CRF-R1 antagonist alone produced no significant change in eIPSCs during LT withdrawal (CP154156 (300 nM): 4.95 ± 1.98% increase relative to baseline; t_(3)_= 0.6470, p = 0.2819). Thus, during LT-withdrawal, CRF-R1 did not seem to be tonically activated and some other functional adaptation must emerge in the weeks following chronic ethanol exposure that limits the ability of CRF, ethanol, and PKC-to facilitate GABAergic neurotransmission.

**Figure 4.**
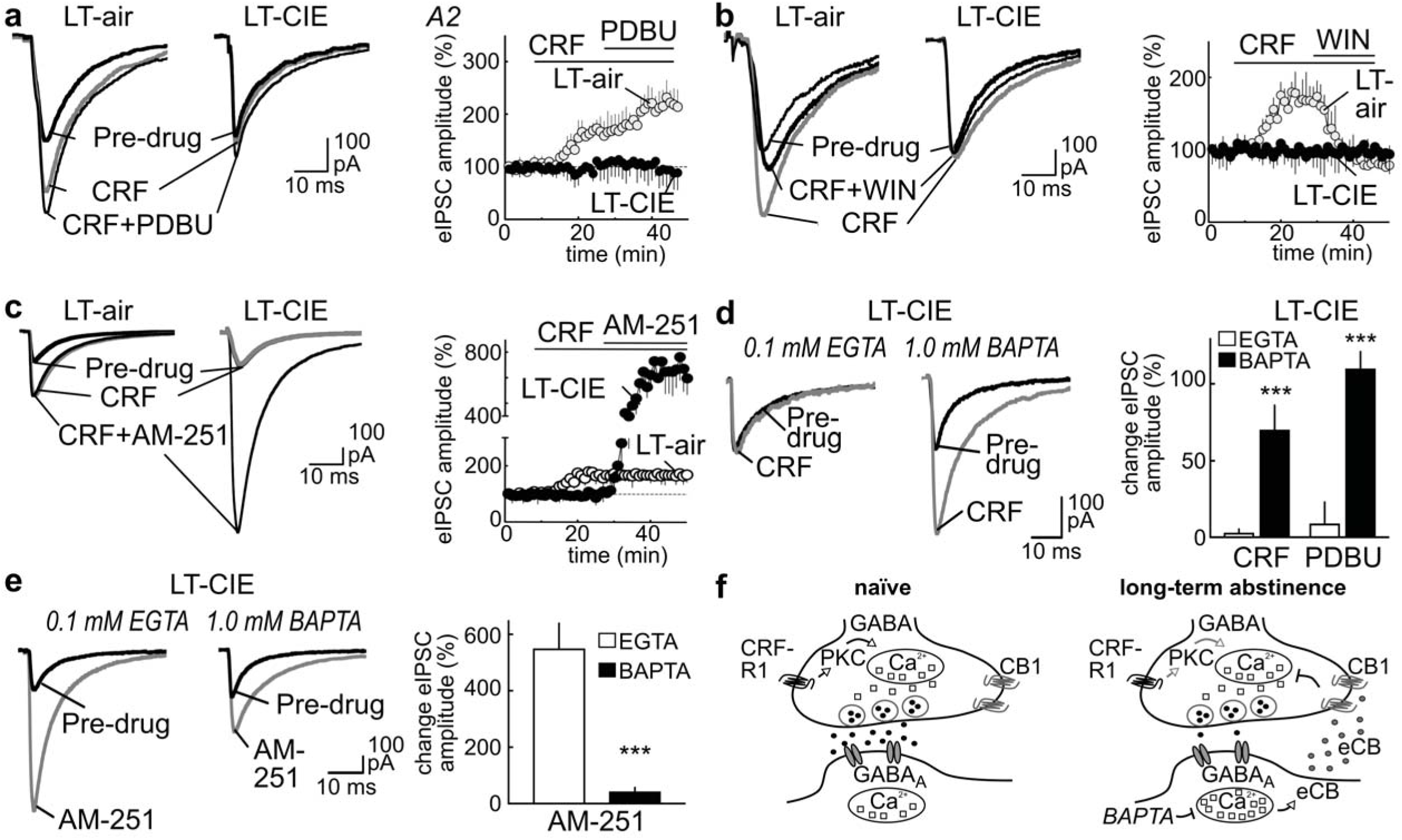
The reduction in CRF-R1 activity during LT withdrawal from CIE is associated with enhanced CB1 receptor mediated inhibition. (**a**) Sample eIPSCs traces and summary data showing reduced responding to CRF (300 nM) and PDBU (1 µM) in neurons from CIE treated mice. (**b,c**) Sample eIPSCs traces and summary data showing LT-CIE cells are (b) less sensitive to CRF (300 nM) and the CB1 agonist WIN-55212 (WIN; 1 µM) but (c) more sensitive to the CB1 antagonist AM251 (1 µM). In (b), note the WIN-related reversal of the CRF potentiation in the air-controls. (**d,e**) Sample eIPSCs traces and summary data illustrating that enhanced calcium buffering (1 mM BAPTA vs 0.1 mM EGTA) in postsynaptic LT-CIE cells rescues the (d) CRF (300 nM) and PdBU (1mM) mediated increase in eIPSC amplitude (Two-way ANOVA, ^***^p < 0.0001 for effect of treatment (internal chelator)), as well as reduces the sensitivity to the (e) CB1 receptor antagonist AM-251 (^***^p < 0.0001). (**f**) Schematic of proposed mechanism of CRF regulation of GABA release onto VTA dopamine neurons during LT withdrawal following CIE. *Left*, in controls, presynaptic CRF-R1 facilitates the release of GABA onto GABA_A_ receptors via PKC mediated recruitment of intracellular calcium. During LT withdrawal from CIE, enhancement of endocannabinoid (eCB) inhibition opposes the CRF action. Chelation of intracellular calcium with BAPTA in the dopamine neuron reduces eCB synthesis, reducing the CB1-mediated suppression of the presynaptic CRF dependent mechanism. (a-d) Number of cells and mice per LT-group: (a) air 8/4, CIE 8/5; (b) air 8/4, CIE 8/5; (c) air 8/4, CIE 8/5;(d) for both EGTA and BAPTA: CRF 7/6, PdBU 6/4; (e) for both EGTA and BAPTA: AM251 6/4. All timelines show eIPSC amplitude % of baseline, mean ± SEM.

We and others have shown that activation of CB1 receptors can suppress VTA GABA release (Riegel and Lupica, 2004; Szabo *et al*, 2014), and in amygdala neurons this reverses the facilitation produced by acute exposure to ethanol (Roberto *et al*, 2010a). Consistent with these reports, we noted that application of the CB1 agonist WIN55212 (1µM) alone in slices from LT-air mice significantly decreased eIPSCs (32.8% ± 4.97% decrease relative to baseline; t_(4)_ = 4.085, p = 0.0075). The same concentrations of WIN55212 reversed the CRF (300 nM) potentiation of eIPSCs (Figure 4b; pre-CRF vs CRF+WIN: t_(4)_ = 0.25465, p = 0.4701). In contrast, in slices from LT-CIE mice, neither CRF nor co-application of WIN55212 with CRF measurably changed IPSC amplitudes (Figure 4b; CRF, t_(4)_ = 0.2314, p = 0.3138; WIN, t_(5)_ = 0.2834, p = 0.3807).

To determine if the diminished actions of CRF and WIN55212 in LT-CIE mice reflected an enhancement of endocannabinoid CB1 receptor inhibition, we measured the effects of CB1R antagonist AM251 (1 µM) on eIPSCs. In control LT-air tissue, AM251 produced no change in eIPSC amplitudes when applied after CRF (Figure 4c; t_(6)_ = 0.2128, p = 0.3781), suggesting the CRF-potentiation of eIPSC amplitude was not restricted by inhibitory tone on CB1 receptors under control conditions. This finding with VTA GABA_A_ IPSCs is consistent with our previous published findings on GABA_B_ IPSCs, which also typically do not show CB1 receptor tone (Riegel and Lupica, 2004). In tissue from LT-CIE mice, as noted above, CRF applied alone produced no change in IPSC amplitude (Figure 4c; t_(6)_ = 0.2128, p = 0.3781). However, application of the CB1 antagonist AM-251 in the presence of CRF produced a robust enhancement of eIPSC amplitude (Figure 4c; t_(6)_ = 32.01, p < 0.0001). This suggests that LT withdrawal from CIE treatment is associated with a functional enhancement of CB1 inhibition on GABA synapses of VTA DA neurons. This CB1 mediated tone blocks subsequent modulation of GABA release by CRF or ethanol and may explain the reduction in the basal frequency of sIPSCs observed in Figure 3d.

As endocannabinoid synthesis requires increases in intracellular calcium in the postsynaptic neuron (Wilson and Nicoll, 2002), we tested whether chelating intracellular calcium post-synaptically in the recorded DA neuron would restore CRF and PdBU sensitivity in LT-CIE tissue. Under normal recording conditions (0.1 mM EGTA) in LT-CIE tissue, CRF and PdBU produced minimal changes in eIPSC amplitude (Figure 4d). In contrast, when recording with an internal solution containing the fast calcium chelator BAPTA (1 mM), both CRF and PdBU enhanced eIPSC amplitude (Figure 4d). A two-way ANOVA that revealed a significant effect of treatment (internal chelator; *F*_(1,16)_ = 51.92, p < 0.0001), but not for drug (*F*_(1,16)_ = 3.892, p = 0.0660) or interaction (*F*_(1,16)_ = 2.126, p = 0.1641). Thus, chelation of intracellular calcium in the postsynaptic neurons of LT-CIE mice restored the ability of bath applied CRF, which acts on the presynaptic CRF-R1, to potentiate GABA_A_ IPSCs.

As a final test that the LT-CIE treatment functionally enhanced endocannabinoid activity at VTA GABA synapses, we measured changes in the amplitude of eIPSCs in the presence of AM-251 under normal recording conditions (0.1 mM EGTA) or altered Ca^2+^ chelation conditions (BAPTA). In the absence of CRF, AM251 again produced a robust increase in eIPSC amplitude in LT-withdrawal (Figure 4e). We observed that the AM251 potentiation of eIPSCs under normal recording conditions was greatly reduced by inclusion of BAPTA in the recording electrode (Figure 4e; *t*_(9)_ = 4.975, p = 0.0008). Taken together, these results indicate that in mice in the weeks following cessation of CIE treatment, the presynaptic CRF-R1 mediated plasticity, typically operative at VTA GABA terminals, is suppressed by a functional enhancement of endocannabinoid activated CB1 receptors (Figure 4f).

## Discussion

We previously reported that in rats activation of presynaptic CRF-R2 facilitated GABA release, and could indirectly reduce glutamate transmission, onto VTA dopamine cells by stimulating presynaptic GABA_B_ receptors (Williams *et al*, 2014). Here we show in mice that CRF increases VTA GABA release via CRF-R1. Acute alcohol produced a similar action (Theile *et al*, 2008; 2011) and we showed that CRF-R1 antagonism did not prevent alcohol-induced GABA release. Short term withdrawal from chronic ethanol exposure did not affect the acute actions of CRF or ethanol, but in subsequent weeks the emergence of endocannabinoid inhibition suppressed the CRF potentiation of GABA release. Such a functional rearrangement of GABAergic inhibition would predictably increase the responsiveness of mesolimbic dopamine neurons to excitatory inputs and this hyper-responsiveness could contribute to long-term vulnerability to relapse following chronic ethanol exposure. In addition to new information about CB1 receptor activation following CIE, this work highlights the differential regulation of GABA between two species and shows an effect of CRF-R1 that has been reported in other regions but not the VTA (Kash *et al*, 2008; Kirby *et al*, 2008; Nie *et al*, 2004; Tan *et al*, 2004).

### Presynaptic CRF-R1 facilitates GABA release

We found that CRF altered the PPR and the frequency of sIPSCs/mIPSCs via GABA_A_ receptors. CRF activated PKC and presynaptic intracellular calcium stores, which serve as reservoirs for release machinery (Llano *et al*, 1994). Calcium entry via voltage-gated calcium channels (VGCC) further augments the release of neurotransmitter (Catterall and Few, 2008). Blockade of VGCCs reduces the frequency of IPSCs (Catterall and Few, 2008) and the CRF facilitation in the VTA. However, only manipulation of ryanodine and IP3 receptors abolished the CRF action, indicating that VGCCs amplify CRF-mediated calcium release from stores (Verkhratsky, 2005). Because calcium stores integrate electrical and chemical signals, these CRF pathways are likely necessary for correctly encoding synaptic plasticity in dopamine neurons (Caillard *et al*, 2000; Verkhratsky, 2005).

CRF G-protein coupled receptors (GPCR) can activate adenylyl cyclase (AC) as well as PKA and PKC, and the intracellular second messenger cAMP, and thus increase levels of free intracellular calcium (for review see (Blank *et al*, 2003;Hauger *et al*, 2009; Riegel and Williams, 2008)). A CRF-R1/PKC action on GABA release also exists in the prefrontal cortex (Tan *et al*, 2004) and the amygdala (Bajo *et al*, 2008). Our findings add the VTA to the growing list of CRF neuromodulations, via both inhibitory (D2 and GABA-B, mGluRs, M1-mAChRs) (Beckstead *et al*, 2009; Riegel and Williams, 2008; Ungless *et al*, 2003; Williams *et al*, 2014) and excitatory (NMDA; HCN) (Korotkova *et al*, 2006; Wanat *et al*, 2008) mechanisms.

### CRF and EtOH regulation of GABA release

Alcohol-induced inhibition has previously been reported in the ventral tegmental area (VTA), substantia nigra, basolateral amygdala (BLA), brainstem, cerebellum and hippocampus of na1ve rodents (Roberto and Varodayan, 2017). Although we did not determine the direct site of action for alcohol, the PKC/PKA/Ca^2+^ signaling pathway is likely involved, as other VTA reports have already shown that acute alcohol increases GABA frequency via 5-HT(2C) receptors, independent of GABA-B and D1-dopamine receptors (Theile *et al*, 2008). Alcohol also increased GABA release in cerebellar interneurons through PKA, PKC and intracellular calcium pathways, and in the amygdala via PKC (Roberto and Varodayan, 2017). In our study, CRF-R1 antagonism did not prevent alcohol-induced GABA release in the VTA of na1ve rats (as previously described in the CeA (Herman *et al*, 2013)), nor did pretreatment with acute CRF yield additive increases with ethanol in na1ve rats. This suggests a convergence of the alcohol and CRF signaling system downstream of the CRF-R1 receptor. Future work must determine if alcohol and CRF actions in the VTA shared a common PKC mechanism (Bajo *et al*, 2008).

### Changes following CIE

Following a history of alcohol dependence and withdrawal, CRF expression is upregulated long-term in multiple brain areas for a period that outlast the brief heightened hypothalamic-pituitary-adrenal axis activation (Becker, 2017). We found that basal GABA release was enhanced after brief withdrawal from chronic ethanol exposure, compared to age matched air-control mice, suggesting greater local inhibition of VTA neurons in alcohol-dependent mice. This may be an action of ethanol common to both the VTA and CeA, as CeA showed similar responses in intoxicated rodents immediately following CIE (Roberto *et al*, 2004; 2010b). However, regional specificity of ethanol’s long-term effects on GABA release exist, since 24 hrs after CIE, GABA release in the dorsal raphe was reduced, but more sensitized to alcohol (Lowery-Gionta *et al*, 2014). Ethanol-induced GABA release in air control mice was consistent with published VTA and amygdala studies (Theile *et al*, 2008; 2011; Varodayan *et al*, 2017) and similar to CRF-induced GABA release. Our observation that acute CRF and alcohol stimulated GABA release in alcohol-dependent mice to a similar magnitude as in na1ve rats, indicates a lack of functional tolerance to acute alcohol or CRF actions at VTA GABAergic synapses. A similar lack of tolerance to alcohol was reported in the CeA of ethanol intoxicated animals immediately after exposure to CIE treatment (5-7 weeks) (Roberto *et al*, 2004). This also means that the previously reported changes in CRF-R1 synthesis, expression and internalization in other regions were presumably either minimal in the VTA or were resolved after 3 days of withdrawal (Reyes *et al*, 2006; Sommer *et al*, 2008).

Following protracted withdrawal, this scenario reversed: basal GABA release normalized in the VTA of alcohol-dependent mice. Moreover, acute CRF or alcohol-induced GABA release was suppressed in the VTA of alcohol-dependent mice relative to air controls, indicating an emergent adaptation in VTA GABAergic synapses to protracted withdrawal. Besides CRF-R1, other G protein-coupled receptors (GPCRs) have been implicated in alcohol-induced CeA GABA release, including the type 1 cannabinoid receptor (CB1) (Roberto and Varodayan, 2017). We found that co-application of a CB1 agonist reversed the acute CRF-induced GABA release in air-control mice, suggesting opposing actions of these two neuromodulators at VTA GABA terminals under baseline conditions. In contrast, CRF no longer induced GABA release, nor did ethanol or the PKC activator PdBU, after long-term withdrawal from CIE. The CB1 receptor agonist also did not alter GABA release after long-term withdrawal from CIE. Application of an antagonist for CB1 receptors produced robust increases in GABA responses after long-term withdrawal, which was not seen in matched air controls. These findings suggest the emergence of an endocannabinoid tone during protracted withdrawal from CIE, given that: 1) chelation of calcium with BAPTA in the postsynaptic membrane reduced the actions of the CB1 antagonist on GABA IPSCs and restored the CRF-induced GABA release in a manner that is conventional for endocannabinoids (Wilson and Nicoll, 2002), 2) the CB1 agonist was ineffective in altering GABA release, and 3) a CB1 antagonist robustly potentiated GABA release in the presence of CRF.

These results coincide with known actions of repeated exposure to ethanol on CB1 receptor inhibition, evoking a transient downregulation of CB1 followed by a long-term upregulation including increased levels of endogenous cannabinoids (Basavarajappa *et al*, 1998; Hungund and Basavarajappa, 2000; Mitrirattanakul *et al*, 2007; Rimondini *et al*, 2002) and studies showing CB1 knockout animals are less sensitive to acute actions of ethanol-associated adaptations (DePoy *et al*, 2013; Lopez-Moreno *et al*, 2012). A more complex role of CB1 receptor inhibition in alcohol-associated changes in the CeA has also been reported (Varodayan *et al*, 2015). Future studies should identify which endocannabinoid mediates this action, whether changes in presynaptic GABA responses induced by acute alcohol are also restored after calcium chelation, and to what extent the function of other GPCRs are impacted.

## Functional implications

Stress plays a pivotal role in alcohol misuse and the magnitude of CRF release during this period determines subsequent stress responsiveness to incentivize excessive drinking (Becker, 2017). During the weeks or months following the binge-like exposure to ethanol associated with the CIE model (Lowery-Gionta *et al*, 2012) the repeated transient homeostatic alterations in CRF signaling eventually stabilize at a new allostatic state (Becker, 2017; Roberto and Varodayan, 2017). The neuroregulation of GABAergic plasticity in the VTA is crucial for burst firing in dopaminergic neurons (Lobb and Paladini, 2010; Paladini *et al*, 1999) and shifts in an enduring manner in the weeks following CIE. The ability of CRF-R1 to potentiate GABA release is suppressed and superseded by a dominating CB1 receptor inhibition of GABA release, which would predictably favor VTA output (Lodge and Grace, 2005). In non-dependent animals, the ethanol enhancement of VTA GABA release onto inhibitory GABA_A_ receptors is typically sufficient to overcome the direct stimulatory effect of ethanol on DA neuron activity (Theile *et al*, 2008; 2011). After repeated alcohol misuse and dependence, a problematic imbalance arises, potentially contributing to sensitizing the VTA to excitatory stimuli thus contributing to the relapse of alcohol seeking.

## Acknowledgements

NIDA grants RO1-DA033342 (A.R.), R37AA009986 (J.W.), U01 AA014095 (H.B.) and a pilot grant from the Alcohol Research Center (ARC; P50-AA10761). We thank Mr. Buchta and Mrs. Pavlinchak for their helpful comments.

